# *NLGN3* autism variants have distinct functional impact on synapses and sleep behavior in *Drosophila*

**DOI:** 10.64898/2026.03.26.714389

**Authors:** Rebekah E. Townsley, Jonathan C. Andrews, Saurabh Srivastav, Sharayu V. Jangam, Shabab B. Hannan, Oguz Kanca, Shinya Yamamoto, Michael F. Wangler

**Affiliations:** Department of Molecular and Human Genetics, Baylor College of Medicine, Houston, Texas, USA; Jan and Dan Duncan Neurological Research Institute, Texas Children’s Hospital, Houston, Texas, USA; Department of Neurology, Baylor College of Medicine, Houston, Texas, USA

## Abstract

*Neuroligin-3 (NLGN3)* was first identified as a risk gene associated with autism spectrum disorder (ASD). The initial variant, p.R451C, associating *NLGN3* with ASD has been heavily investigated, yet little is known about the functional consequences of other *NLGN3* variants. Furthermore, while most of the identified variants are present in males with maternally inherited variants from unaffected mothers, several *de novo* variants were observed in females, suggesting a possible functional difference between *de novo* and maternally inherited variants. To address the functional consequences of *NLGN3* variants *in vivo*, we generated transgenic *Drosophila* models corresponding to one *de novo* variant (p.R175W) identified in one female proband, and two maternally inherited variants (p.R451C and p.R597W) identified in male probands. In *Drosophila*, loss of the fly homolog, *Nlg3,* altered sleep patterns, synaptic architecture, and vesicle dynamics, which were rescued by the expression of the human *NLGN3^Ref^* allele. When comparing the variants, the *de novo* p.R175W variant and the maternally inherited p.R451C variant altered synapse morphology and sleep patterns, with minimal effects on vesicle dynamics, and the p.R597W variant altered sleep and vesicle dynamics with minimal impact on synapse morphology. Using overexpression models, human *NLGN3^Ref^* altered sleep patterns and synaptic morphology. Moreover, the p.R175W variant exacerbated sleep phenotypes, and the p.R175W and p.R451C variants exacerbated synapse morphology phenotypes. Together, our findings suggest that *de novo NLGN3* variants identified in females are likely gain-of-function, while maternally inherited variants have mixed loss-and gain-of-function effects. Moreover, the location of the variants may contribute to the distinct functional differences we observed. Some *NLGN3* variants disrupt synaptic development, while other variants alter synaptic function, suggesting that *NLGN3* variants have differential effects. These functional differences may provide insight into the heterogeneity of individuals with ASD.

**Author Summary:** Autism spectrum disorder (ASD) is a common neurodevelopmental disorder. Mutations in the *Neuroligin-3 (NLGN3)* gene are associated with ASD but very few of these mutations have been characterized in animal models. Most of these mutations affect male individuals who maternally inherited their genetic mutation; however, more rarely female individuals may present with a genetic mutation that was not identified in either of the parents. Here, we utilized the fruit fly model to investigate how three different mutations, one mutation identified in a female and two mutations identified in males, affect the fly’s behavior and synapse development. We identified altered sleep patterns in some of our mutants which is consistent with sleep disturbances being highly comorbid with ASD. Additionally, we identified alterations in synapse development and function which is consistent with the role of *NLGN3* in synapse formation and maturation. Together, our findings support that *NLGN3* is important for regulating the synapse and mutations in this gene can alter its function. However, different mutations can have differential effects. This demonstrates the need to assess multiple variants simultaneously because each variant may have distinct functional significances.

## Introduction

Autism spectrum disorder (ASD) is a highly heritable complex neurodevelopmental disorder characterized by deficits in social communication and interaction, and restricted, repetitive behaviors [1]. It affects approximately 1 in 31 children in the United States annually, with boys diagnosed 3.4 times more than girls [2]. Additionally, about 74% of individuals with ASD have at least one other co-occurring neurological or psychiatric disorder, including intellectual disability, attention-deficit hyperactivity disorder, anxiety, and sleep disorders [3–6]. Although numerous ASD susceptibility genes have been identified, many converge on pathways regulating neuronal communication and brain development, including neurite outgrowth and synaptic function [7–10].

The *Neuroligin* (*NLGN*) gene family was among the first genes associated with ASD [11]. NLGNs are a family of postsynaptic cell adhesion molecules that regulate synapse formation and maturation with their binding partner, Neurexin [12–15]. Their intracellular domain contains postsynaptic density 95/discs large/zona occluden-1 (PDZ) and gephyrin binding motifs that indirectly recruit GABAergic and glutamatergic receptors to the postsynaptic membrane [16,17]. This signaling is essential for establishing synaptic contacts and maintaining synaptic plasticity, both crucial for proper brain function. Humans have five *NLGN* genes that are preferentially expressed at either the excitatory or inhibitory synapses; however, only *NLGN3* is uniquely localized to both [18–21]. *NLGN3,* located on Xq13.1, was first associated with ASD when a maternally inherited p.R451C variant was identified in two brothers with ASD [11,22]. Since then, its functional impact has been heavily investigated in mouse knock-in and human cell models [23–30]. Although most studies support a gain-of-function (GoF) mechanism, emerging evidence suggests a cell type–dependent effect: some brain regions show GoF while others show loss-of-function (LoF) properties [23,26,27,31]. Beyond the p.R451C variant, at least 23 additional *NLGN3* variants have been associated with ASD, most inherited maternally [32,33]. While the p.R451C variant has been well studied, the functional impact of other ASD-associated *NLGN3* variants remains less understood.

Since *NLGN3* is located on the X-chromosome, mechanistic and functional differences may exist between affected males and females. Notably, all maternally inherited variants are observed in affected males with unaffected maternal carriers, whereas *de novo* variants affect males and females. These inheritance patterns suggest that *NLGN3* variants may act through multiple mechanisms, with maternally inherited variants functioning differently from *de novo* variants in females. To determine if there are distinct mechanisms between affected males and females, we modeled variants of interest *in vivo*.

Our lab previously used the *Drosophila melanogaster* model organism to screen 79 *de novo* ASD variants in 74 genes and identified that 38% of these variants cause functional changes in *Drosophila* [34]. Using a humanization strategy, we can determine whether variants act in a LoF, GoF, or in a complex context-dependent manner [34–38]. With a more simplified neuronal architecture, the *Drosophila* neuromuscular junction (NMJ) also provides functional insight into genes regulating synapse development and plasticity [39]*. Drosophila* has four *Neuroligin* (*Nlg*) genes; *Drosophila Nlg3* is homologous to human *NLGN3* [40–43]. Furthermore, *Drosophila Nlg* genes are known to alter synaptic development at the NMJ. *Nlg1*, *Nlg2,* and *Nlg4* mutants reduce bouton number while *Nlg3* mutants increase bouton number, and all *Nlg* mutants impair synaptic transmission [40,42–46]. Like its mammalian counterpart, *Nlg3* is also the only member that is expressed in both excitatory and inhibitory synapses [47,48]. While loss of *Nlg3* has been characterized in knockout models, we and others have developed transgenic toolkits to rapidly assess the functional impact of human missense variants [40,47]. In this study, we utilized *Drosophila* to investigate whether human NLGN3 can functionally replace the fly protein and assess the functional impact of three ASD-associated *NLGN3* variants: two maternally inherited missense variants and one female *de novo* missense variant. We investigated their impact on *Drosophila* behavior, synapse morphology, and vesicle dynamics through humanization and overexpression strategies. These findings help elucidate the functional impact of *NLGN3* variants and assess whether maternally inherited and *de novo* variants act through distinct mechanisms.

## Results

### Selection of variants

*NLGN3* was associated with ASD in 2003 when a maternally inherited missense variant (p.R451C) was observed in two brothers [11]. Since then, at least 23 *NLGN3* missense variants have been identified in individuals with ASD, many of which remain functionally uncharacterized [32,33]. Of these, 5 are *de novo*, 13 are maternally inherited, and 5 are of unknown inheritance. Notably, two *de novo* variants were observed in females, while all remaining variants—including maternally inherited and remaining *de novo* cases—were observed in males (S1 Table). There are several X-linked genes where the genetic mechanism differs between maternally inherited and *de novo* variants. For example, *de novo* loss of function *MECP2* variants cause Rett syndrome primarily in females whereas *de novo* duplication and triplication cause *MECP2* duplication syndrome primarily in males [49,50]. Additionally, *de novo DDX3X* variants in females cause X-linked dominant intellectual disability, but maternally inherited variants cause X-linked recessive intellectual disability in males [51,52]. Most *NLGN3* variants affect males, and these variants are typically inherited from unaffected female carriers; however, more rarely, affected females exhibit *de novo* variants. We, therefore, hypothesized that *de novo* variants in females confer a GoF mechanism whereas maternally inherited variants in males have mixed LoF and GoF effects.

**Table S1:** Identified 23 ASD-associated *NLGN3* variants and used bioinformatic programs to prioritize variants of interest.

| Variant<br>(BC051715.1<br>transcript) | cDNA | Inheritance<br>Pattern | Biological<br>Sex | Allele<br>Frequency | CADD | PolyPhen2 | SIFT | Publication |
| --- | --- | --- | --- | --- | --- | --- | --- | --- |
| p.P120L | c.359 C>T | *De Novo* | Male | 8.26E-07 | 28.7 | 0.802 | 0 | Laurie et al<br>(2025) |
| **p.R175W** | **c.523 C>T** | ***De Novo*** | **Female** | **1.65E-05** | **30** | **0.982** | **0** | **Iossifov et al. (2014)** |
| p.G406S | c.1216 G>A | *De Novo* | Female | 8.25E-07 | 26.7 | 0.023 | 0.01 | Xu et al.<br>(2014) |
| p.R640C | c.1918C>T | *De Novo* | Male | 3.30E-06 | 25.9 | 0.761 | 0 | ClinVar<br>(2024) |
| p.L721R | c.2162 T>G | *De Novo* | Male | Absent | 17.95 | 0.063 | 0.19 | Seth et al.<br>(2023) |
| p.G188R | c.562G>A | Maternal | Male | Absent | 33 | 0.995 | 0 | Qin et al.<br>(2025) |
| p.S249N | c.746G>A | Maternal | Male | Absent | 24.2 | 0.001 | 0.03 | ClinVar<br>(2021) |
| p.I252F | c.754A>T | Maternal | Male | Absent | 28.5 | 0.946 | 0 | ClinVar<br>(2022) |
| p.V326M | c.976 G>A | Maternal | Male | 3.31E-06 | 24.2 | 0.102 | 0.05 | Jiang et al.<br>(2013) |
| p.V341A | c.1022 T>C | Maternal | Male | 2.31E-05 | 26.1 | 0.698 | 0 | Yu et al.<br>(2013) |
| **p.R451C** | **c.1351 C>T** | **Maternal** | **Males (x2)** | **Absent** | **33** | **1** | **0** | **Jamain et al. (2003)** |
| p.P514S | c.1540 C>T | Maternal | Males (x2) | Absent | 26 | 0.97 | 0 | Quartier et al. (2019) |
| p.M538V | c.1612A>G | Maternal | Male (x3) | Absent | 27 | 0.998 | 0 | ClinVar<br>(2021) |
| p.T542I | c.1625C>T | Maternal | Male | Absent | 31 | 0.962 | 0 | ClinVar<br>(2024) |
| p.D594N | c.1780 G>A | Maternal | Male | Absent | 27.5 | 0.793 | 0 | Sedlackova et al. (2024) |
| **p.R597W** | **c.1789 C>T** | **Maternal** | **Males (x3)** | **8.29E-07** | **28.8** | **1** | **0** | **Redin et al. (2014), Quartier et al. (2019)** |
| p.T632A | c.1894 A>G | Maternal | Males (x2) | 3.83E-03 | 20.7 | 0 | 0.08 | Blasi et al. (2006) |
| p.R777W | c.2329C>T | Maternal | Male | Absent | 24.8 | 1 | 0 | ClinVar (2023) |
| p.E139K | c.415G>A | Unknown | Male | 8.26E-07 | 32 | 1 | 0.02 | ClinVar (2025) |
| p.Y147H | c.439 T>C | Unknown | Male | Absent | 28.3 | 1 | 0 | Greco et al. (2025) |
| p.N216S | c.647A>G | Unknown | Unknown | Absent | 25 | 0.965 | 0 | ClinVar (2021) |
| p.A442T | c.1324G>A | Unknown | Unknown | Absent | 32 | 0.958 | 0 | ClinVar (2022) |
| p.H761R | c.2282A>G | Unknown | Unknown | 7.52E-06 | 18.56 | 0.28 | 0.02 | ClinVar (2024) |

This table summarizes all ASD-associated *NLGN3* variants published in SFARI or ClinVar databases with the year the variant was submitted. Variants selected for functional analysis are bolded. The recurrence of a variant within a family are noted in the parentheses by the biological sex. Allele frequency was obtained from gnomAD. CADD: combined annotation dependent depletion; PolyPhen2: polymorphism phenotyping 2; SIFT: sorting intolerant from tolerant

To assess whether specific variants may act through distinct pathogenic mechanisms, we queried SFARI and ClinVar databases for ASD-associated *NLGN3* missense variants [32,33]. We identified 13 variants from SFARI and 10 variants from ClinVar that were associated with ASD. Using *in silico* prediction tools, we selected the top three variants with the highest combined annotation dependent depletion (CADD) score where the biological sex and inheritance pattern were known: two maternally inherited (p.R451C and p.R597W) and one *de novo* (p.R175W) variant present in a female (S1 Table). Notably, p.R451C and p.R597W variants have been previously studied *in vivo* or *in vitro,* respectively [11,53,54].

### Bioinformatic evaluation of ASD-associated *NLGN3* variants

The first missense allele (p. R175W) was identified as a *de novo* variant in a heterozygous female, diagnosed with ASD and intellectual disability (Fig. 1A) [55]. The second variant (p.R451C) was maternally inherited in two hemizygous brothers diagnosed with ASD (Fig. 1A) [11]. The third variant (p.R597W) was maternally inherited in three hemizygous males all diagnosed with ASD, intellectual disability, and sleep disorders (Fig. 1A) [53,54]. While the last two variants were inherited, none of the maternal carriers were affected. All three variants are localized to the cholinesterase-like (Che-like) domain, the primary extracellular domain of NLGN3, which is predicted to be highly intolerant to change by MetaDome analysis, suggesting a potential hotspot for pathogenic variants (Fig. 1B) [56]. This domain interacts with Neurexin to regulate synapse maturation, suggesting that mutations in this domain may disrupt synaptic function. To assess the pathogenic effect of these three variants, we utilized *in silico* prediction tools. *In silico* analysis predicted that the p.R451C and p.R597W variants are likely pathogenic, while the p.R175W variant yielded inconsistent results (S2 Table). We also used MutPred2 to predict specific molecular alterations that may be causative of disease [57]. Mutpred2 predicted distinct molecular mechanisms for the p.R175W variant while the other two variants had similar molecular mechanisms. Specifically, the p.R175W variant is predicted to become more rigid and less regulated, suggesting it may alter its interaction with other binding partners. While the other two variants are predicted to alter the transmembrane, suggesting it may be misfolded or mislocalized (S2 Table). Lastly, all three variants affect highly conserved residues. The p.R451 and p.R597 residues are conserved across all vertebrate and invertebrate species analyzed as well as across all *NLGN* family members; however, the p.R175 residue is only conserved in vertebrates and mildly conserved among *NLGN* paralogs (Fig. 1C and Figure S1A). These findings suggest that these residues are critical for the function of NLGN3; however, they may have different functional consequences. To study this, we used the *Drosophila melanogaster* model organism.

**Fig. 1:**
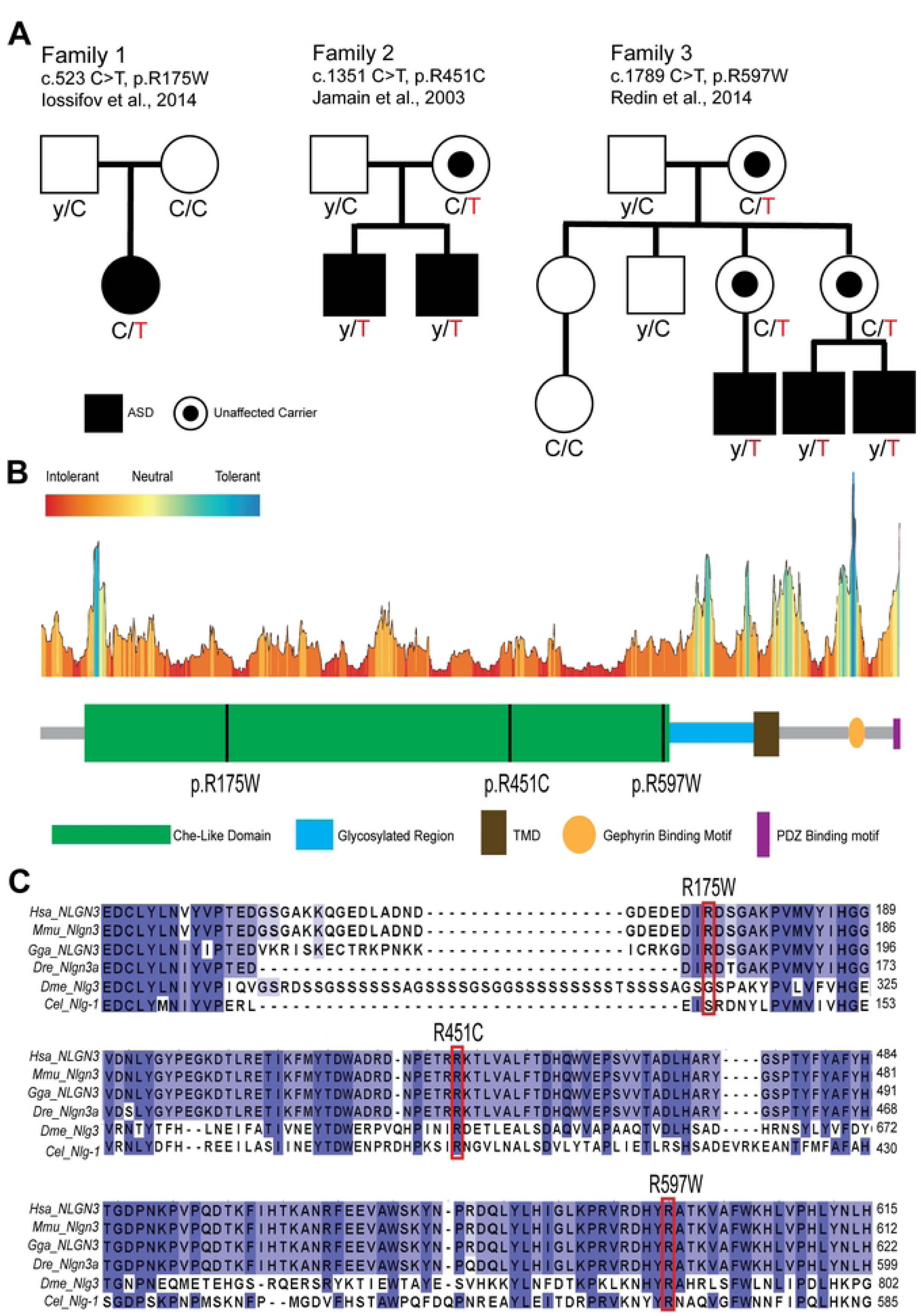
Pedigree and conservation analysis. **A)** Family pedigrees. Open squares or circles represent unaffected males or females, respectively. Closed squares or circles represents affected males or females, respectively. A filled circle inside an open circle represents unaffected female carriers. **B)** Functional domains of NLGN3 and variants’ location using MetaDome analysis. Functional domains of NLGN3 include cholinesterase-like domain (Che-like), glycosylated region, transmembrane domain (TMD), Gephyrin binding motif, and PSD-95/DLG1/ZO-1 binding motif (PDZ motif). **C)** Evolutionary conservation of *NLGN3* variants. Human (BC051715.1), mouse (Q8BYM5), chicken (A0A8V0Z5J3), zebrafish (D0EVX1), *Drosophila* (Q9VIC6), and C. elegans (Q9XTG1) were used.

**S2 Table:**
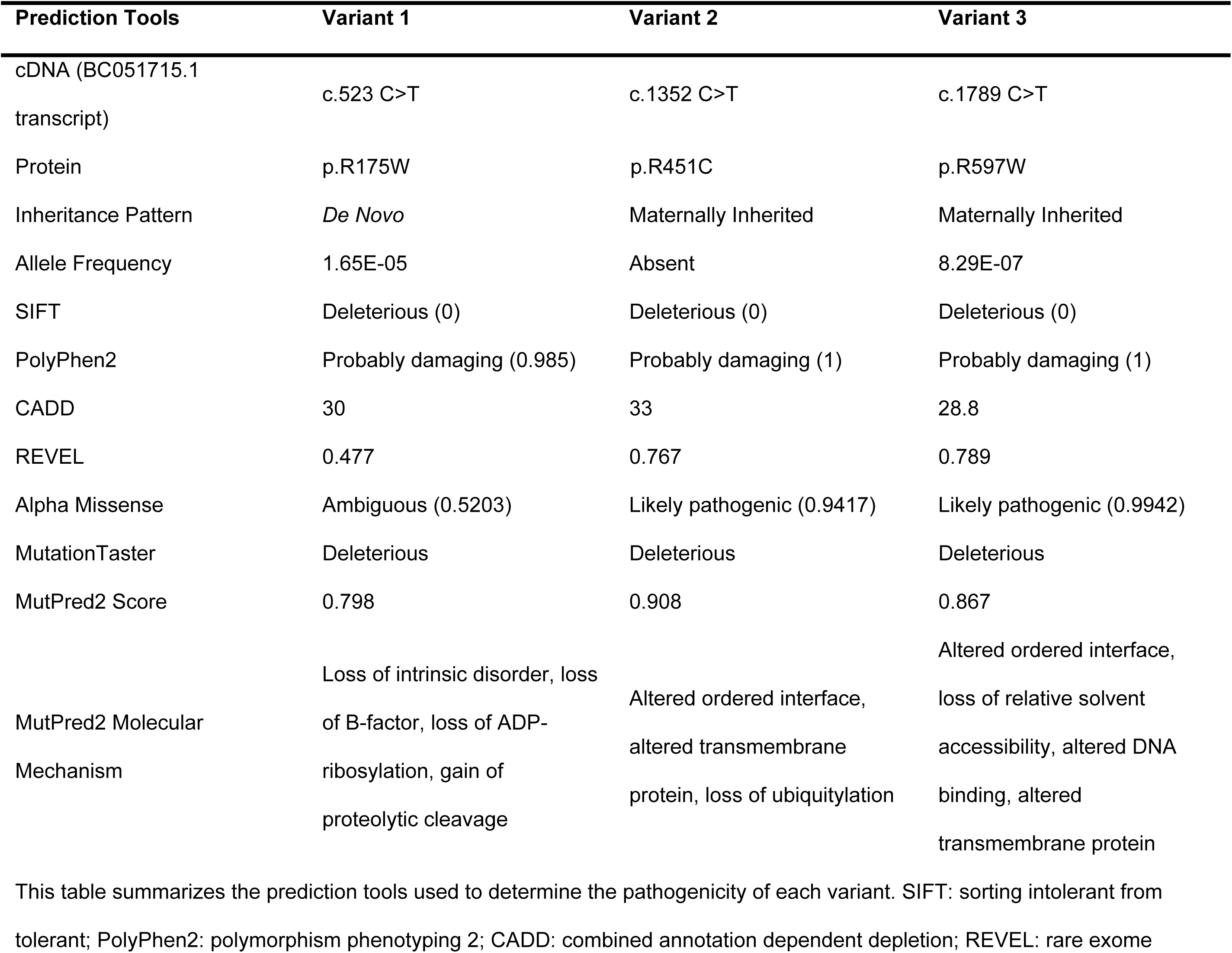
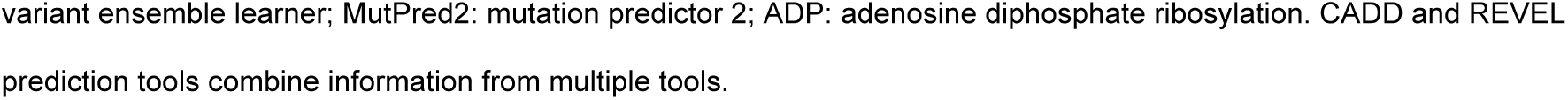
Summary of in silico prediction tools used to determine the pathogenic effect of three ASD-associated NLGN3 variants.

### *Drosophila Nlg3* is expressed in neurons

*Drosophila* has four *Nlg* family genes: *Nlg1, Nlg2, Nlg3,* and *Nlg4* [40–43]. *Nlg3* shows the highest homology to human *NLGN3* (DIOPT score = 12, 32% identity and 46% similarity) [58]. To study the functional impact of patient-derived mutations, we used a previously generated *Nlg3*-specific trojan GAL4 (TG4) allele [34,59]. The *Nlg3^TG4^* allele contains a splice donor, T2A ribosomal skipping sequence, GAL4 transcriptional activator, and a polyA signal, and it is inserted between exons 3 and 4 (Fig. 2A) [59]. This insertion truncates the endogenous protein and expresses a GAL4 transcriptional activator in the same spatial and temporal manner. The GAL4 protein can drive the transcription of cDNAs that are cloned downstream of an upstream activation sequence (UAS). This allele can be used to detect the expression pattern of fly *Nlg3* and express human *NLGN3* cDNA in the expression domain of *Nlg3*. *NLGN3* is primarily expressed in neurons but has been identified in astrocytes and plays a role in neuron-glia interaction [60]. To assess the lineage trace expression pattern of fly *Nlg3*, we crossed *Nlg3^TG4^* with a G-Trace reporter and stained adult brains with neuronal marker Elav and glial marker Repo [61]. Similar to humans, *Nlg3* is primarily expressed in neurons and sparsely expressed in glia throughout development (Fig. 2B-I and Figure S1B-G).

**Fig. 2:**
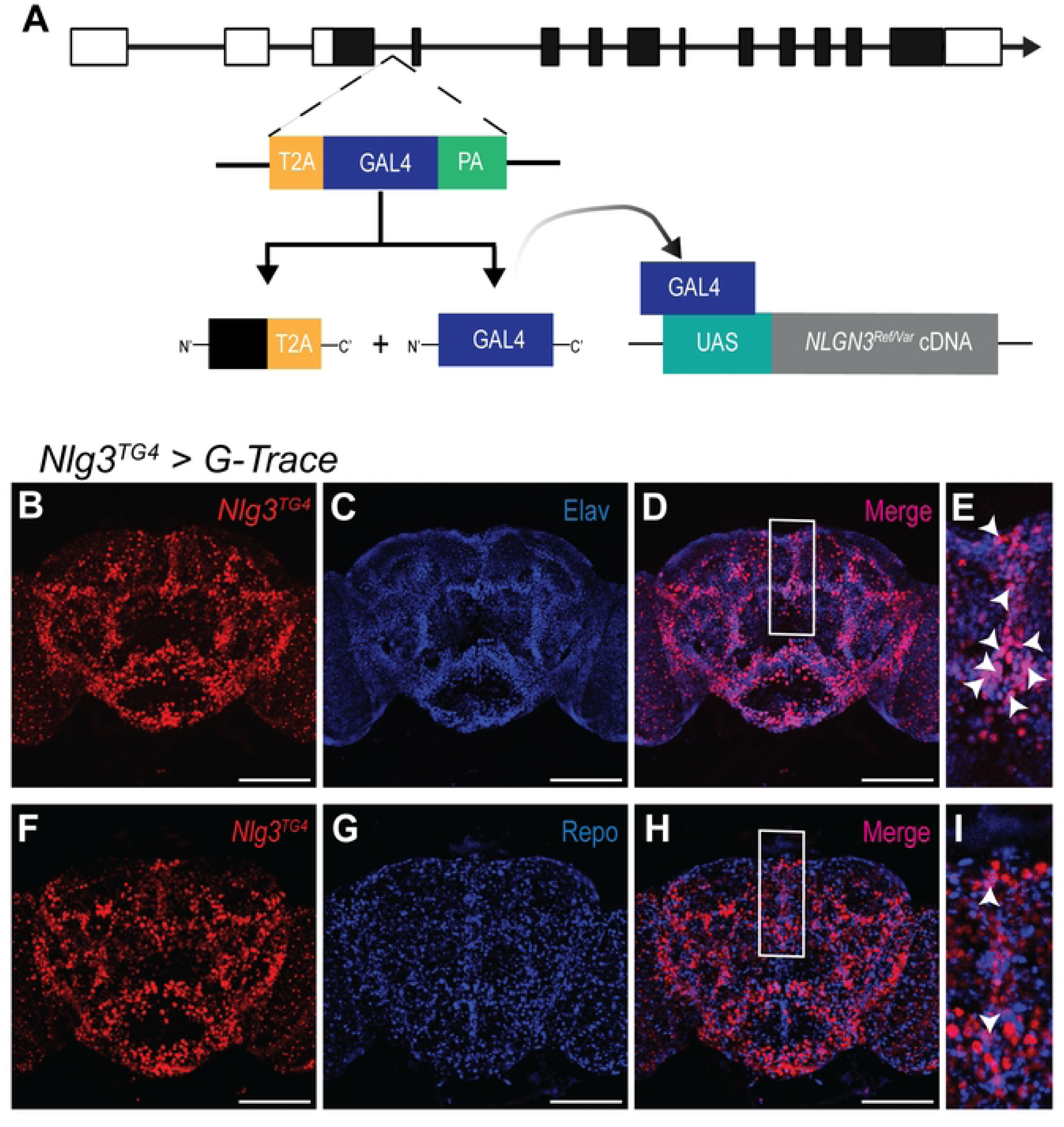
Humanization strategy enables spatiotemporal control of *Nlg3* expression. **A)** The *Nlg3* specific trojan GAL4 (*Nlg3^TG4^*) allele truncates the fly protein but produces a GAL4 transcriptional activator in the same spatiotemporal pattern as the endogenous fly gene. The GAL4 protein can drive the expression of UAS-*NLGN3* constructs or a fluorescent reporter. **B, F)** *Nlg3^TG4^* driving a G-trace reporter line was used to identify the current expression pattern of *Drosophila Nlg3*. **C)** anti-Elav staining labels neurons. **D, E)** Merging the expression of *Nlg3* and Elav positive cells identifies that *Nlg3* is expressed in neurons. **G)** anti-Repo staining labels glial cells. **H, I)** Merging *Nlg3* and Repo positive cells shows some overlap between *Nlg3* and glia suggesting *Nlg3* is sparsely expressed in glia. White arrowheads indicate colocalization. Scale bar: 100 μm.

### *NLGN3* alters locomotion and sleep behaviors

Although the *Nlg3^TG4^* allele is predicted to function as a strong LoF allele, it has not been fully characterized. To confirm this, we first quantified the total mRNA expression in adult heads and compared it to a known null allele (*Nlg3^Null^*) [47]. Using primers downstream of the insertion site, we observed that the *Nlg3^TG4/Null^* flies have 85% and *Nlg3^Null/Null^* flies have 99% decrease in *Nlg3* mRNA levels; however, both mutants are viable (Fig. 3A). To determine if loss of *Nlg3* leads to neuronal phenotypes such as a shortened lifespan, we recorded longevity in *Nlg3* mutants. Interestingly, control and *Nlg3^Null/Null^* flies had a median survival rate of 46 days, but the *Nlg3^TG4/Null^* flies had a median survival rate of 57 days (Fig. 3B). This could be attributed to the *Nlg3^TG4^* being a hypomorph rather than an amorph or differences in genetic backgrounds. To further assess if the *Nlg3^TG4^* allele shows phenotypic similarities to the *Nlg3^Null^* alleles, we next assessed locomotor activity. Loss of *Nlg3* is known to cause deficits in locomotion [40,47]. To measure motor activity, we recorded basal activity of *Nlg3^TG4/Null^* and *Nlg3^Null/Null^* flies using the *Drosophila* activity monitor [62]. Consistent with previous literature, we observed a significant decrease in locomotion in both of these allelic combinations, suggesting that *Nlg3^TG4^* is likely functioning as a LoF allele [40,47]. However, the expression of the human *NLGN3^Ref^* and missense variants in a *Nlg3^TG4/Null^* background failed to rescue this activity defect (Fig. 3C & Figure S2A). These findings were recapitulated when we measured the amount of time the fly spent moving within a test chamber (Figure S2B). When we overexpressed *NLGN3^Ref^* using the neuronal driver Elav-GAL4, we observed a decrease in total activity, but the activity/waking minute was not significant, suggesting that motor function is not impaired (Figure S2C-D). Together these results suggests that fly *Nlg3* may play a role in basal locomotion that cannot be rescued by human *NLGN3*.

**Fig. 3:**
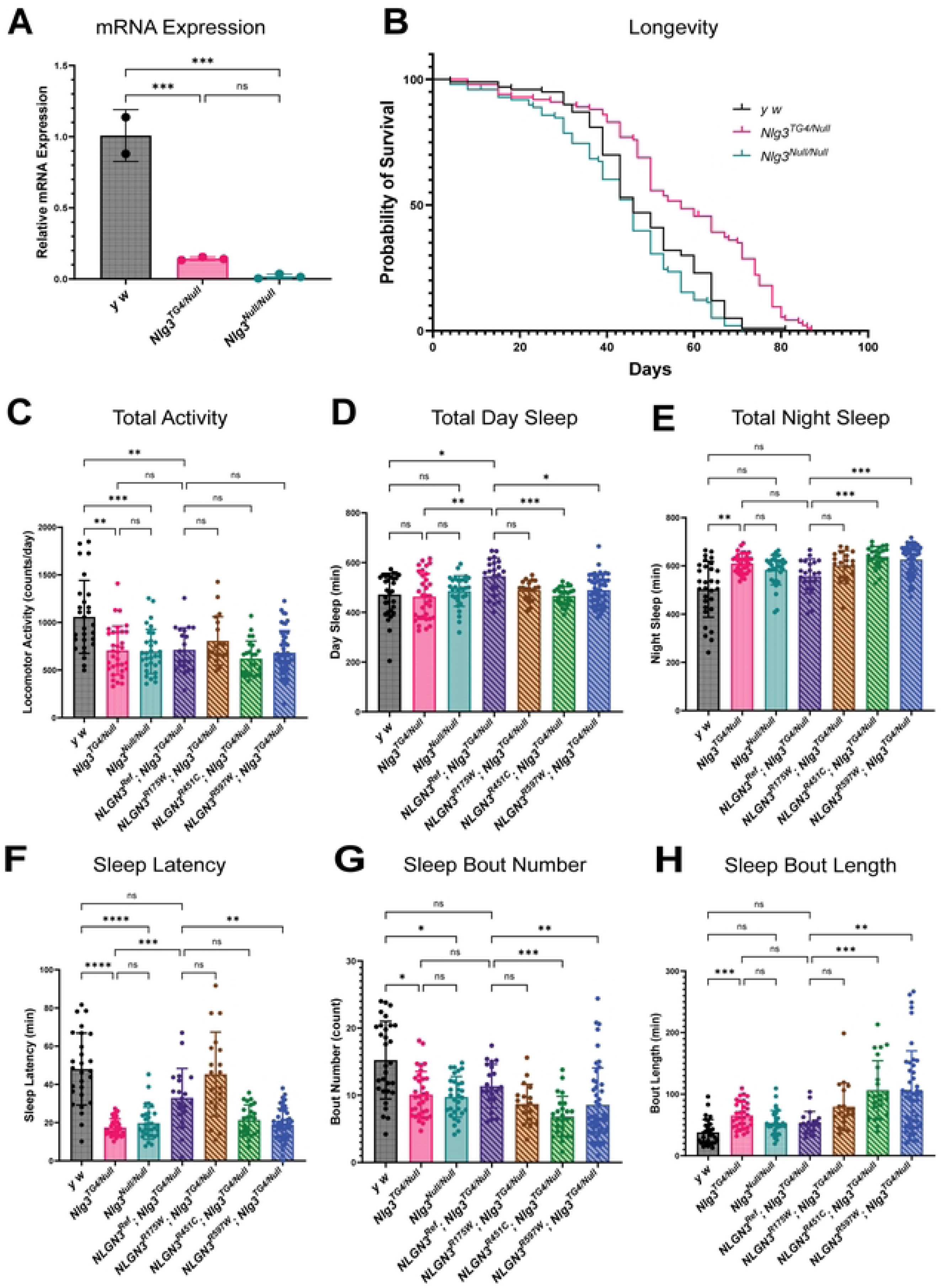
*NLGN3* variants have altered sleep patterns. **A)** Relative mRNA expression in *Nlg3^TG4/Null^* and *Nlg3^Null/Null^* flies (2-3 biological replicates, n=20 heads per biological replicate, 3 technical replicates; One-way ANOVA with Sidak’s multiple comparison test, ns *P* > 0.05, \*\*\**P* ≤ 0.001). **B)** Survival curves of *y^1^ w\** (controls), *Nlg3^TG4/Null^* and *Nlg3^Null/Null^* flies. *Nlg3^TG4/Null^* mutants have increased lifespan compared to controls (χ^2^ = 26) while *Nlg3^Null/Null^* mutants have reduced lifespan compared to controls (χ^2^ = 5.18) (n=100 adult flies per genotype, Mantel Cox Logrank test). **C)** Quantification of total locomotor activity (n≥19 flies; Welch’s ANOVA with Dunnett’s T3 multiple comparison test, ns *P* > 0.05, ** *P* ≤0.01, *** *P* ≤0.001). **D)** Quantification of sleep duration during lights on (n≥19 flies). **E)** Quantification of sleep duration during lights off (n≥19 flies). **F)** Quantification of sleep latency (n≥19 flies). **G)** Quantification of sleep bout number during lights off (n≥19 flies). **H)** Quantification of sleep bout length during lights off (n≥19 flies). Kruskal Wallis with Dunn’s multiple comparison test was used for D-H; ns *P* > 0.05, * *P* ≤0.05 ** *P* ≤0.01, *** *P* ≤0.001).

Since ASD and sleep are highly comorbid and one of the variants was associated with sleep disorders in multiple individuals, we next assessed whether *Nlg3* is important for sleep function [53,54]. Previous rodent models have implicated *NLGN3* in sleep disruptions, particularly alterations in the sleep/wake cycle [63,64]. We found that loss of *Nlg3* does not affect the duration of daytime sleep, but nighttime sleep was increased. Interestingly, the expression of the *NLGN3^Ref^* in a *Nlg3^TG4/Null^* background increased total daytime sleep while partially rescuing nighttime sleep defects. The expression of all three variants in a *Nlg3^TG4/Null^* background failed to rescue the duration of nighttime sleep (Fig. 2D-E). Loss of *Nlg3* also drastically reduced sleep latency and increased sleep consolidation while expression of *NLGN3^Ref^* in a *Nlg3^TG4/Null^* background was able to partially rescue both phenotypes. The expression of the *NLGN3^R451C^* and *NLGN3^R597W^* variants in a *Nlg3^TG4/Null^* background further exacerbated sleep latency and sleep consolidation defects while the expression of the *NLGN3^R175W^* variant was able to fully rescue sleep latency and had no additional effect on sleep consolidation (Fig. 2F-H). Furthermore, overexpressing *NLGN3^Ref^* using the neuronal driver Elav-GAL4 increased the duration of day and night sleep, decreased sleep latency and the number of sleep bouts, and increased the length of each sleep bout. However, overexpressing the *NLGN3^R175W^* variant with Elav-GAL4 driver further exacerbated many of these phenotypes indicating a GoF mechanism (Figure S2E-I). Together, our findings suggest that *NLGN3* variants can cause neuronal phenotypes.

### *NLGN3* alters synaptic morphology

Since *NLGN3* regulates synapse formation and maturation in multiple species, we hypothesized that *NLGN3* variants may affect synapse morphology and activity, driving these behavioral phenotypes in individuals with ASD. To evaluate synaptic architecture, we first used the third instar *Drosophila* larval NMJ, which is primarily glutamatergic and has large, easily visualized boutons [39]. Loss of *Nlg3* is known to increase the number of type 1B boutons and reduce bouton size while branch number remains unchanged [40]. Co-staining the NMJ with horse radish peroxidase (HRP), a neuronal membrane marker, and discs large (DLG), a postsynaptic subsynaptic reticulum marker, we recapitulated these findings with our LoF mutants (Fig. 4A-C). The expression of *NLGN3^Ref^* and *NLGN3^R597W^* in a *Nlg3^TG4/Null^* background rescued this bouton overgrowth phenotype whereas *NLGN3^R175W^* and *NLGN3^R451C^* variants showed persistent bouton overgrowth, similar to the LoF mutants, suggesting that these variants are unable to rescue bouton growth (Fig. 4A-H). Consistent with previous findings, NMJ branch number was unchanged (Figure S3). Although *NLGN3^Ref^* rescued bouton number, bouton size remained reduced, indicating that some functions of *Drosophila Nlg3* cannot be rescued by *NLGN3* (Fig. 4I).

**Fig. 4:**
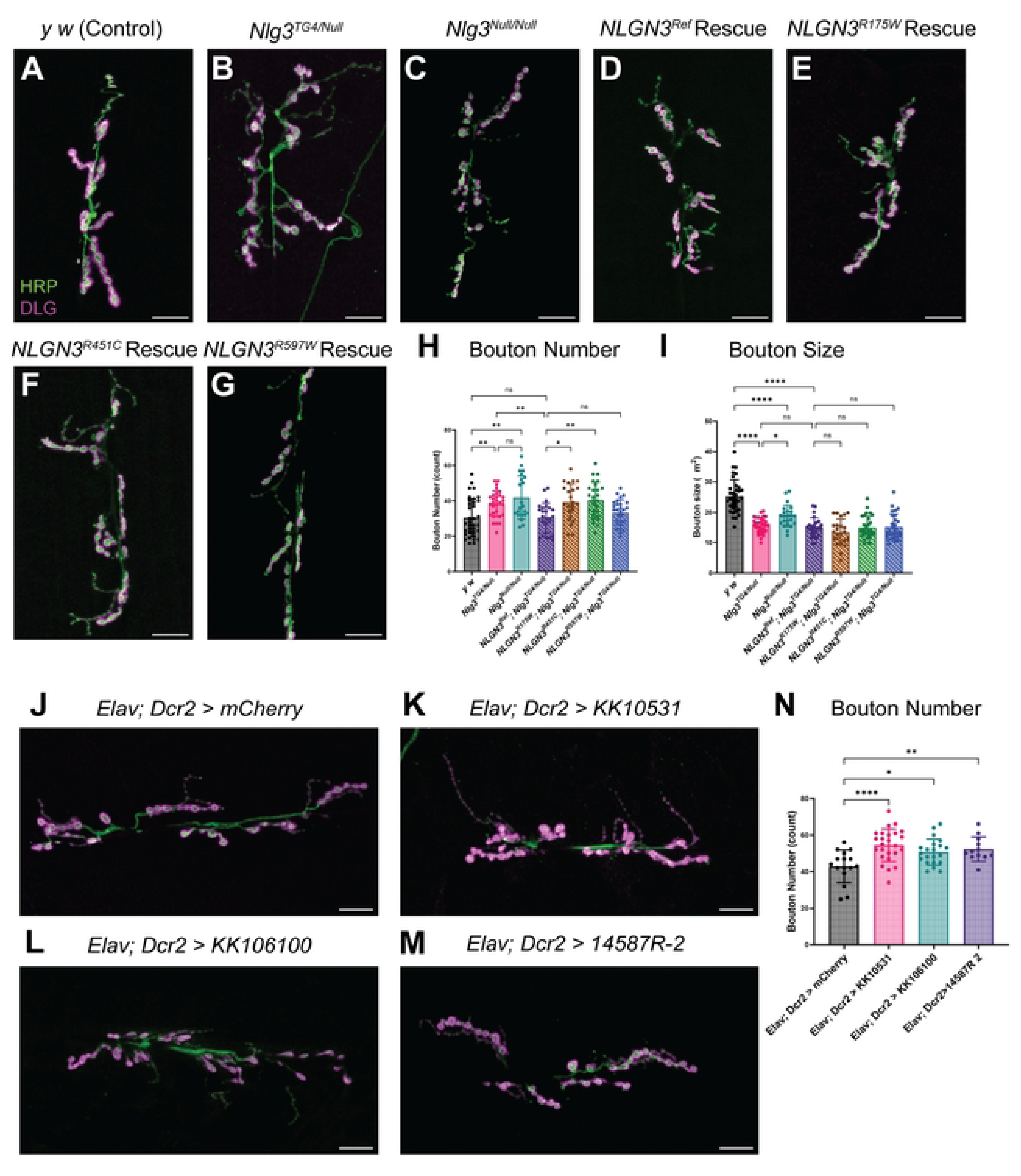
*NLGN3* variants alter synaptic architecture at the neuromuscular junction. **A-G)** Representative images of type 1B boutons at muscle 6/7 of abdominal segment A4 from *y*^1^ *w*\**, Nlg3^TG4/Null^, Nlg3^Null/Null^, NLGN3^Ref^; Nlg3^TG4/Null^, NLGN3^R175W^; Nlg3^TG4/Null^, NLGN3^R451C^; Nlg3^TG4/Null^*, and *NLGN3^R597W^; Nlg3^TG4/Null^* larvae. Neuromuscular junctions (NMJ) were co-stained with anti-HRP, a pan neuronal marker, and anti-DLG, a postsynaptic marker. **H)** Quantification of the number of boutons at each NMJ (n≥22 larvae/genotype, Welch’s ANOVA with Dunnett’s T3 multiple comparison test; ns *P* > 0.05, * *P* ≤0.05, **** *P* ≤ 0.0001. **I)** Quantification of bouton size (n≥22 larvae/ genotype, Welch’s ANOVA with Dunnett’s T3 multiple comparison test; ns *P* > 0.05, * *P* ≤ 0.05, **** *P* ≤ 0.0001). **J-M)** Representative images of type 1B boutons in *Nlg3* knockdown models (*LacZ, KK10531, KK106100, 14587R-2*) using a neuronal driver (Elav-GAL4). NMJs were co-stained with anti-HRP and anti-DLG. **N)** Quantification of bouton number in *Nlg3* knockdown models (n≥12 larvae, One-way ANOVA with Sidak’s multiple comparison test; ns *P* > 0.05, * *P* ≤0.05, ** *P* ≤0.01, **** P ≤0.0001. Scale bar: 20 μm.

*NLGN3* is primarily expressed in neurons; however, some evidence suggests fly *Nlg3* may be expressed in low levels at the larval muscle wall [40]. However, immunofluorescence did not detect *Nlg3* expression at the muscle wall. Thus, it remains unclear which tissue is relevant for bouton growth phenotype. To assess tissue-specific effects, we used overexpression and knockdown models. Overexpressing *NLGN3^Ref^* and *NLGN3^R597W^* with *Nlg3^TG4^* driver in *Nlg3^TG4/+^* larvae did not alter bouton number; however, overexpressing *NLGN3^R175W^* and *NLGN3^R451C^* variants increased bouton number (Figure S4A-B). Similar phenotypes were observed when overexpressing human *NLGN3* cDNAs with neuronal driver, Elav-GAL4, and motor neuron driver, D42-GAL4 (Figure S4C-F). In contrast, muscle specific overexpression (Mef2-GAL4) reduced bouton number even in *NLGN3^Ref^* expressing animals (Figure S4G-H). Using knockdown models, we identified that the neuronal knockdown of *Nlg3* using Elav-GAL4 increased bouton number in three independent RNAi lines, resembling LoF mutants (Fig. 4J-N). This suggests that the loss of *Nlg3* in neurons is sufficient to cause bouton overgrowth and the expression of *NLGN3* variants in the neurons is likely causing the phenotypes we observed.

Since our behavioral assays were performed in adult flies, we next assessed if morphological defects observed in larvae could also be observed in adult flies. To determine this, we examined NMJ morphology in 7-day-old flies. Staining the abdominal body wall with cysteine string protein (CSP), a presynaptic marker, we observed an increase in the number of boutons in *Nlg3* mutants (Figure S5A-C). The expression of *NLGN3^Ref^* and *NLGN3^R597W^* in a *Nlg3^TG4/Null^* background was able to rescue this bouton phenotype (Figure S5D and S5G). However, similar to the larval NMJ, the expression of the *NLGN3^R175W^* and *NLGN3^R451C^* variants both failed to rescue bouton overgrowth (Figure S5E-H). These results indicate that *Nlg3* in *Drosophila* regulates bouton growth throughout development and some *NLGN3* variants impair bouton regulation.

### *NLGN3* variants alter vesicle dynamics

To further examine *NLGN3’s* role in synapse maturation, we next assessed the morphology of the active zone protein, Bruchpilot (BRP). Co-staining the third instar larval NMJ with BRP and HRP, revealed a significant increase in the number of BRP puncta in the LoF mutants when normalized to the bouton size (Fig. 5A-C). The expression of the *NLGN3^Ref^* in a *Nlg3^TG4/Null^* background was able to fully rescue this observation (Fig. 5D). Similar to the overgrown bouton phenotype observed earlier, we saw an increase in BRP puncta in animals rescued with either *NLGN3^R175W^* or *NLGN3^R451C^* variants (Fig. 5E-F). However, the *NLGN3^R597W^* variant did not disrupt BRP puncta (Fig. 5G-H). The core region of the Che-like domain is important for interacting with Neurexin; therefore, variants (p.R175W and p.R451C) within this hotspot may impair synapse development by disrupting transsynaptic interactions. Together, *NLGN3* seems to be important not only for facilitating synaptic growth but also for active zone regulation.

**Fig. 5:**
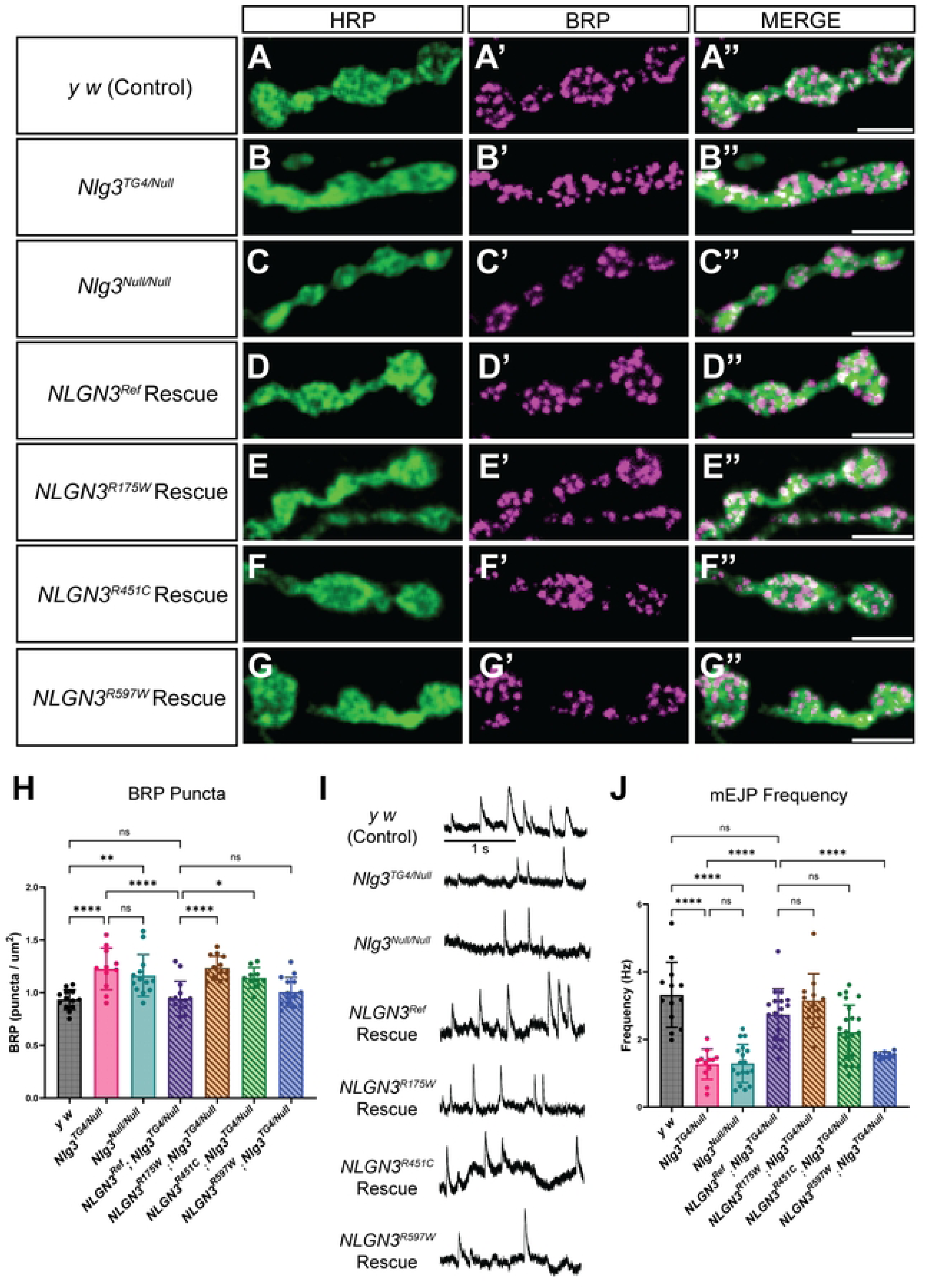
*NLGN3* variants impair vesicle dynamics. **A-G)** Representative images of type 1B boutons at muscle 6/7 of abdominal segment A4 from *y^1^ w*, Nlg3^TG4/Null^, Nlg3^Null/Null^, NLGN3^Ref^*; *Nlg3^TG4/Null^, NLGN3^R175W^; Nlg3^TG4/Null^, NLGN3^R451C^; Nlg3^TG4/Null^, and NLGN3^R597W^; Nlg3^TG4/Null^* larvae. Boutons were co-stained with HRP, a pan-neuronal marker, and BRP, an active zone marker. Scale bar: 5 μm. **H)** Quantification of the number of BRP puncta normalized to the size of each bouton (N≥10 larvae/genotype, n=6 boutons/larvae). One-way ANOVA with Sidak’s multiple comparison test; ns *P* > 0.05, * *P* ≤ 0.05, ** *P* ≤ 0.01, **** *P* ≤ 0.0001. **I)** Representative miniature excitatory junction potentials (mEJP) traces at muscle 6 of abdominal segments A3 and A4 from *y^1^ w*, Nlg3^TG4/Null^, Nlg3^Null/Null^, NLGN3^Ref^; Nlg3^TG4/Null^, NLGN3^R175W^; Nlg3^TG4/Null^, NLGN3^R451C^_;_ Nlg3^TG4/Null^*, and *NLGN3^R597W^; Nlg3^TG4/Null^* larvae. Each spike corresponds to one mEJP event. Scale bar: 1 second. **J)** Quantification of the frequency of mEJP (n≥9). Welch’s ANOVA with Dunnett T3 multiple comparison test; ns *P* > 0.05, **** *P* ≤ 0.0001.

Neurotransmitters are released at the site of active zones; however, not all active zones are functional. To determine if these active zones are functional, we recorded miniature excitatory junction potentials (mEJPs) at the third instar larval NMJ over two minutes. Consistent with previous research, LoF mutants showed a decrease in the mEJP frequency that was fully rescued when *NLGN3^Ref^* was expressed in a *Nlg3^TG4/Null^* background [40]. Surprisingly, the expression of the *NLGN3^R175W^* and *NLGN3^R451C^* variants fully or partially rescued this phenotype, respectively; however, the expression of the *NLGN3^R597W^* variant failed to rescue this defect (Fig. 5I-J). Additionally, we did not observe any difference in the mEJP amplitude or kinetics (Figure S6). This suggests that the presence of BRP puncta does not correlate with the ability of the variants to rescue vesicle release. This is likely because not all active zones are fully functional. It is possible that in the p.R175W and p.R451C variants, having an increase in active zones may help restore the threshold needed to maintain the frequency of spontaneous vesicle release.

*Nlg3* has been implicated not only in synapse development but also in synaptic vesicular trafficking [40]. To determine if our missense variants disrupt vesicle trafficking, we used the FM1-43 styryl dye which functions in an activity dependent manner. Similar to previous reports, *Nlg3* LoF mutants had a significant decrease in dye uptake compared to control animals, indicative of an endocytic defect (Figure S7A-C)[40]. The expression of human *NLGN3* reference or variant constructs in a *Nlg3^TG4/Null^* background rescued this dye uptake defect (Figure S7D-H). This suggests that the expression of *NLGN3* reference and variants are sufficient to restore endocytosis. However, the dye distribution within the bouton appeared abnormal in the variants tested. In both the control and *NLGN3^Ref^* rescue animals, the dye appeared as puncta within the boutons; however, in the LoF mutants and variant rescue animals, the majority of the dye surrounded the border of the boutons suggesting that vesicle size and/or morphology may be altered.

## Discussion

Although *NLGN3* is an established ASD susceptibility gene, most variants have yet to be functionally characterized. In this study, we selected three *NLGN3* variants with high CADD scores and modeled them in *Drosophila*. Consistent with previous findings, loss of *Nlg3* reduced locomotor activity, increased the number of type 1B boutons, and decreased frequency of mEJP and FM1-43 dye uptake [40,47]. Additionally, we observed phenotypes that were not previously reported such as decreases in sleep latency and increases in nighttime sleep duration and BRP puncta (Table 1). Based on humanization strategies, we rescued many of these LoF phenotypes using human *NLGN3*, demonstrating that human NLGN3 can functionally replace the *Drosophila* protein in many biological instances (Table 1). We then examined the functional consequences of three ASD-associated variants. We showed that the *NLGN3^R175W^* variant, a *de novo* variant identified in a female with ASD, was also able to rescue sleep latency, frequency of mEJP, and FM1-43 dye uptake phenotypes (Table 1). However, when we overexpressed this variant with neuronal drivers, these animals had exacerbated sleep phenotypes and the boutons were significantly overgrown compared to *NLGN3^Ref^* flies, indicating a GoF mechanism (S3 Table). The *NLGN3^R451C^* variant exhibited a complex mechanism and cannot be classified as a simple LoF or GoF allele. When we expressed this variant in a *Nlg3^TG4/Null^* background, it failed to rescue sleep and NMJ morphological defects, suggesting it has some LoF properties (Table 1). Yet, when we overexpressed this variant, the boutons were significantly overgrown compared to *NLGN3^Ref^*, suggesting it also possesses some GoF properties (S3 Table). Overall, our study points towards a complex mechanism that is likely context dependent. Lastly, the *NLGN3^R597W^* variant failed to rescue sleep deficits and frequency of mEJP; however, it was able to restore NMJ morphology and FM1-43 dye uptake (Table 1). Overexpression of this variant with neuronal drivers, did not produce additional phenotypes, likely conferring a LoF mechanism (S3 Table). Together these results suggest that variants in *NLGN3* can produce a spectrum of functional outcomes, and individual variants contribute to behavioral phenotypes through mechanistically distinct ways.

**Table 1:**
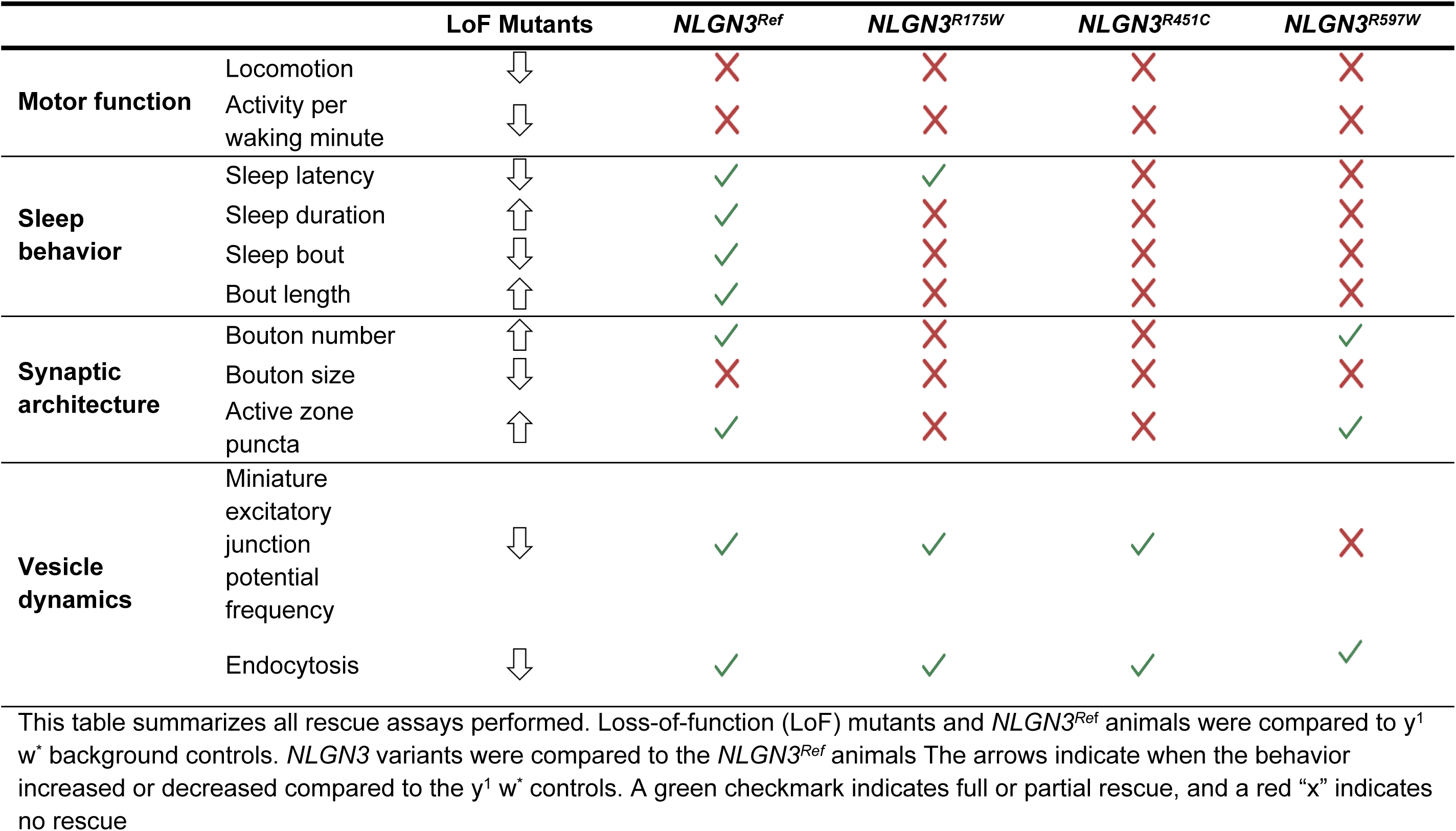
Summary of all rescue assays performed.

**S3 Table:** Summary of overexpression assays performed.

|  |  | *NLGN3<sup>Ref</sup>* | *NLGN3<sup>R175W</sup>* | *NLGN3<sup>R451C</sup>* | *NLGN3<sup>R597W</sup>* |
| --- | --- | --- | --- | --- | --- |
| **Motor function (Neuronal driver)** | Locomotion | ↓ | ↓ | ns | ↓ |
|  | Activity per waking minute | ns | ns | ns | ns |
| **Sleep behavior (Neuronal Driver)** | Sleep latency | ↓ | ns | ns | ns |
|  | Sleep duration | ↑ | ↑ | ns | ↑ |
|  | Sleep bout | ↓ | ↓ | ns | ns |
|  | Bout length | ↑ | ↑ | ns | ns |
| **Synaptic architecture** | Bouton number (*Nlg3<sup>TG4</sup>* driver) | ns | ↑ | ↑ | ns |
|  | Bouton number (neuronal driver) | ns | ↑ | ↑ | ns |
|  | Bouton number (motor neuron driver) | ns | ↑ | ↑ | ns |
|  | Bouton number (muscle driver) | ↓ | ns | ns | ns |

This table summarizes overexpression assays performed. *NLGN3<sup>Ref</sup>* animals were compared to *LacZ* animals. *NLGN3* variants were compared to *NLGN3<sup>Ref</sup>* animals. Arrows indicate whether the behavior increased or decreased from their respective controls. ns = not statistical difference when compared to their respective controls.

The functional investigation of *NLGN3* variants in ASD has been mostly limited to the *NLGN3^R451C^* allele [11]. Although *NLGN3* variants were originally considered GoF, several early stop gain variants have been identified, suggesting that LoF *NLGN3* variants may also contribute to disease phenotypes [65,66]. Numerous *NLGN3^R451C^* mouse knock-in studies have supported a causative role in ASD pathophysiology. Several studies have recapitulated ASD-relevant behaviors including deficits in social interaction, hyperactivity, enhanced repetitive motor routines, and surprisingly, enhanced spatial learning abilities [23,27,30]. Consistent with the hypothesis that ASD may alter the excitation/inhibition balance, the *NLGN3^R451C^* variant increases the strength of inhibitory synaptic transmission in the somatosensory cortex and reduces inhibitory synaptic transmission at synapse formed by Parvalbumin basket cells [23,31]. Other studies have shown it increases the strength of the excitatory synaptic transmission in the hippocampus, yet these effects were not observed in *NLGN3* knockout mice suggesting a GoF mechanism [23,27,31]. Still, others observed an increase in inhibitory synaptic transmission in cholecystokinin positive GABAergic basket cells that was also seen in *NLGN3* knockout mice suggesting a LoF mechanism [31]. Together, these studies suggest that *NLGN3* may induce cell-type and synapse specific changes. In our study, we observed both LoF and GoF properties for the *NLGN3^R451C^* variant. While *NLGN3^R451C^* failed to rescue sleep and NMJ morphological defects, suggesting a LoF phenotype, overexpressing this variant further exacerbated bouton overgrowth, suggesting a GoF phenotype. Our results are consistent with previous studies, further confirming a context-dependent effect. Moreover, *NLGN3* variants affect both males and females, yet there is a male-specific bias. While most of the variants are maternally inherited in affected males, several *de novo* variants in affected females have been observed [55,67]. Overall, this suggests that maternally inherited and *de novo NLGN3* variants likely have distinct functional mechanisms, otherwise, we would expect the maternal carriers to also phenotypically be affected.

Individuals with ASD are more likely to experience sleep deficits. Roughly 50-80% of individuals with ASD will experience sleep disturbances which is two to three times higher than typical developing children [68]. Sleep disturbances can negatively impact the quality of life not only for the child but also their caregivers [69]. Although the relationship between sleep and ASD is relatively unclear, several studies have implicated *NLGN3* in sleep disruptions [63,64]. *NLGN3* knockout and *NLGN3^R451C^* knock-in mice have alterations in spectral power bands during different sleep stages, suggesting that *NLGN3* can affect sleep quality [63,64]. Furthermore, the conditional knockout of *NLGN3* in medial septum neurons in mice causes hyperactivity of GABAergic neurons and results in social memory impairment and sleep disturbances [70]. While *Drosophila Nlg3* has not been implicated in sleep disruptions prior to our study, sleep abnormalities have been reported in *Drosophila Nlg4* mutants [41]. *Drosophila Nlg4* is highly expressed in the large lateral ventral neurons (l-LNvs) and fan shaped bodies, the sleep centers of the fly brain [41]. Loss of *Nlg4* impaired GABA transmission resulting in hyperactivity and increase sleep latency [41]. Together these studies suggest that the excitatory/inhibitory imbalance may contribute to the pathogenesis of both ASD and sleep disturbances.

In the context of sleep, we were particularly interested in one variant, *NLGN3^R597W^*, which was identified in three affected individuals who were diagnosed with ASD and sleep disorders [53,54]. Driven by clinical data, we assessed if *Drosophila Nlg3* may alter sleep behaviors. We identified that loss of *Nlg3* increased nighttime sleep duration and reduced sleep latency, yet these phenotypes were rescued when *NLGN3^Ref^* was expressed, suggesting that *Nlg3* is involved in sleep in flies as well. Furthermore, *NLGN3* variants failed to rescue sleep duration and sleep consolidation phenotypes, confirming that our assays are sensitive enough to detect sleep defects in the *NLGN3^R597W^* variant. While the *NLGN3^R451C^* and *NLGN3^R597W^* variants behaved similarly in the sleep assay, the *NLGN3^R597W^* variant did not have any NMJ morphological defects and failed to rescue the mEJP frequency like the *NLGN3^R451C^* variant. Together this suggests that even the maternally inherited variants may have distinct functional mechanisms.

Although we did not explore the trafficking of *NLGN3* variants, MutPred2 predicted that the *NLGN3^R451C^* and *NLGN3^R597W^* variants will likely impair membrane signaling leading to the mislocalization of the protein. This was confirmed in previous *in vitro* studies that identified the *NLGN3^R451C^* and *NLGN3^R597W^* variants are retained in the endoplasmic reticulum (ER) [23,24,53]. Additionally, MutPred2 predicted that the *NLGN3^R451C^* variant could accumulate misfolded proteins which can lead to cellular stress. This was validated using *in vitro* studies that observed an accumulation of NLGN3 proteins in the ER which activated the unfolded protein response (UPR) leading to a 90% in cell surface protein expression [23,24,53]. However, 10% of the protein is able to be trafficked properly leading to some functional NLGN3 [23]. In contrast, the *NLGN3^R597W^* variant is predicted to alter DNA binding and impair proper posttranslational modification in addition to impairing membrane signaling. *In vitro* assays suggest that this variant is unable to localize properly to the plasma membrane and increases the activation of the UPR [53]. However, additional *in vivo* studies on the localization of this variant are needed. These molecular differences may provide insight into how these variants are altering function. Although the cellular trafficking of the *NLGN3^R175W^* variant has yet to be studied, MutPred2 does not predict mislocalization with this variant but rather alterations in binding interactions. Therefore, the *NLGN3^R597W^* variant may impair synaptic function without altering synapse development; however, other variants may disrupt synaptic architecture by impairing proper binding interactions or by interacting with noncanonical partners [71]. Understanding whether the *NLGN3^R175W^* variant impairs vesicle trafficking or inhibits binding with presynaptic partners can provide mechanistic insight into how different mutations across the protein can differentially affect synapse function.

While we have developed transgenic toolkits to assess the pathogenicity of missense variants in *Drosophila*, our study does have some limitations. Although the expression of human *NLGN3^Ref^* in a *Nlg3^TG4/Null^* background rescued many LoF phenotypes, it failed to rescue basal locomotor activity and only partially rescued sleep defects. Previous studies demonstrated that the short isoform of wild-type fly *Nlg3* produced in neurons is critical for regulating locomotor behavior [45]. It is unclear from our study whether human *NLGN3* can undergo the same cleavage process. One possible explanation is that human *NLGN3* is unable to be cleaved into a short and long isoform, thus locomotion is not restored.

In this study, we assessed synapse morphology at the NMJ. The *Drosophila* NMJ is experimentally more accessible and structurally more simplified compared to the mammalian brain [39]. The stereotyped architecture of the NMJ and its capacity for developmental and functional plasticity, make it a powerful system for probing the synaptic consequences of *NLGN3* variants [39]. In our study, electrophysiological recordings were performed at the muscle. While *NLGN3* is generally considered a postsynaptic membrane protein primarily expressed in neurons, low levels of fly *Nlg3* mRNA have been detected in the muscle tissue, but the corresponding protein was undetectable [40,45]. Therefore, it is unclear whether *NLGN3* is acting as a postsynaptic protein in the muscle, acting as a presynaptic protein in the dendrites of motor neurons, or acting upstream in the central nervous system. We observed that knockdown of fly *Nlg3* using neuronal drivers was sufficient to alter bouton morphology. However, other studies demonstrated that the expression of *Nlg3* in the muscle and not neurons is able to rescue bouton morphology [40]. Thus, it is possible that the expression of *Nlg3* in both the central nervous system and muscle may regulate NMJ development [44]. Further studies are needed to tease apart this circuit.

In summary, we investigated the functional impact of three ASD-associated *NLGN3* variants using *Drosophila*. We identified functional differences between *de novo* variants observed in females and maternally inherited variants observed in males. We showed that a *de novo* variant that has not been functionally studied previously is likely a GoF allele, whereas maternally inherited variants are likely more complex than originally proposed. Additionally, we observed functional differences between the two maternally inherited variants where one variant effects synapse development and the other effects synaptic function. From our study, we propose *de novo* variants in females are GoF, maternally inherited variants within the core region of the Che-like domain regulate synapse development, and maternally inherited variants that are distal to the binding sites regulate synaptic function. Further studies are needed to mechanistically understand these different proposed models. However, this study provides new insight into the functional differences between *NLGN3* variants and how these functional differences may confer a spectrum of phenotypes seen in individuals with *NLGN3* variants.

## Materials/Subjects and Methods

### Bioinformatic methods

ASD-associated *NLGN3* variants were identified through the Simons Foundation Autism Research Initiative (SFARI) and ClinVar databases [32,33]. To prioritize variants, we utilized *in silico* prediction models such as gnomAD, SIFT, PolyPhen2, CADD, REVEL, Alpha Missense, MutationTaster, and MutPred2 [57,72–80]. Clustal Omega was used to align multiple sequences obtained from UniProt and the Jalview software was used to visualize sequence alignment [81–84]. MetaDome was used to visualize pathogenic hotspots [56].

### *Drosophila* stocks and husbandry

Fly stocks were maintained at room temperature, unless otherwise specified. Experiments were conducted in a temperature and humidity-controlled incubator at 25°C, 50% humidity, and a 12-hour light/dark cycle. Flies were reared on standard fly food (water, yeast, soy flour, cornmeal, agar, corn syrup, and propionic acid). The Canton-S and *p{UAS-lacZ}* flies were provided by Hugo Bellen (Baylor College of Medicine, Texas, USA) and the *Nlg3^Def1^*(*Nlg3^Null^*) flies were provided by Anne Simon (University of Western Ontario, Canada). The remaining fly stocks were obtained from Bloomington *Drosophila* stock center (Indiana University, Indiana, USA), Vienna *Drosophila* research center (Vienna, Austria), and Kyoto *Drosophila* stock center (Kyoto Institute of Technology, Kyoto, Japan): *y^1^ M{vas-int.Dm}ZH-2A w*; PBac{y[+]-attP-3B}VK00037* (BDSC #24872), *y^1^ w*; Mi{Trojan-GAL4.2}Nlg3MI00445-TG4.2 (Nlg3^TG4^*) (BDSC #76134), *w*; P{GawB}D42* (BDSC #8816), *P{GawB}elav[C155]* (BDSC #458), *P{GAL4-Mef2.R}3* (BDSC #27390), *w*; P{UAS-RedStinger}4, P{UAS-FLP.D}JD1, P{Ubip63E(FRT.STOP)Stinger}9F6/CyO* (G-Trace) (BDSC #28280), *y^1^ sc* v^1^ sev^21^ P{VALIUM20-mCherry.RNAi}attP2* (BDSC #35785), *P{GawB}elav[C155]; P{UAS-Dcr-2.D}2* (BDSC #25750), *P{KK106100}VIE-260B* (VDRC: v100376), *P{KK110531}VIE-260B* (VDRC: v102055), *P{NIG.14597R}2* (NIG-Fly: 14597R-2)

### Human *NLGN3* transgenic line generation

Human *NLGN3* reference clone (BC051715.1 transcript in pDonr221 plasmid) was obtained from the Kenneth Scott human cDNA collection (Department of Molecular and Human Genetics, Baylor College of Medicine). Variant constructs (p.R174W, p.R451C, and p.R597W) were generated using NEBase Changer Q5 site-directed mutagenesis kit (New England Biolabs Inc #E0554S) and verified by Sanger sequencing. Reference and variant coding sequences (CDS) were subcloned into the p.UASg-HA.attB vector using LR clonase enzymes (Thermo Fisher Scientific, #12538120)[85]. Verified plasmids were microinjected into *vas-phiC31; VK0037* embryos [86]. All constructs were injected into the VK37 docking site to avoid positional effects. Transgenic male flies were crossed to *y^1^ w\** virgin female flies, and successful integration was confirmed by selecting for the mini white gene.

### RNA extraction and Real-time quantitative PCR

Total RNA was extracted from heads of 20 adult flies per genotype using *Quick*-RNA Miniprep Plus kit (Zymo Research #R1057) followed by cDNA synthesis using iScript cDNA Synthesis kit (Bio-Rad #1708890). cDNA concentrations were normalized followed by quantitative PCR experiments using iQ SYBR Green Supermix (Bio-Rad #1708880) on a CFX Opus 96 Real-Time PCR system (Bio-Rad, California, USA). Gene expression was normalized to the *rp49* gene. Primers were designed with PrimerBlast and purified by high-performance liquid chromatography. Primer sequences are as follows:

*rp49* Fwd 5’-GCTAAGCTGTCGCACAAATG-3’
*rp49* Rev 5’-GTTCGATCCGTAACCGATGT-3’
*Nlg3* Fwd 5’-GCAATATGGAATCGAGGCGG-3’
*Nlg3* Rev 5’-AATCACCGAATCGCGTCTG-3’

### Longevity Assay

Eclosed flies were separated by sex and genotype and maintained at 25°C. 10 flies of the same sex and genotype were group housed together. Flies were transferred into a new vial every 3-4 days and survival was determined. A Log-rank Mantel Cox test was used to determine significance.

### *Drosophila* sleep and activity

Five-to-seven-day old male flies were placed in *Drosophila* Activity Monitors (DAM, Trikinetics, Massachusetts, USA) and housed at 25°C, 50% humidity on a 12-hour light/dark cycle for seven days. Inactive periods longer than five minutes were classified as sleep. Total activity and sleep behaviors were analyzed using MATLAB SleepMat software [87]. Sleep latency was defined as time from lights off to sleep onset. Sleep consolidation is composed of the number of sleep bouts and the average sleep bout length.

Locomotor behavior was performed as previously described [88]. Socially naive males were collected by isolating pupae in 16 × 100 polystyrene vials with 1 mL of fly food. Locomotion was assessed in a custom made 6 well acrylic [89]. One 6-to-10-day old mutant or rescue male fly was introduced into the chamber well through aspiration. Locomotion was recorded with a Basler 1920UM, 1.9MP, 165FPS, USB3 Monochromatic camera and the BASLER Pylon module, with an adjusted capturer rate of 33 frames per second. Videos were compiled using a custom MATLAB script. Fly positions were tracked with CalTech Flytracker [90].

### Brain immunostaining

*Nlg3^TG4^* was crossed with *w*; P{UAS-RedStinger}4, P{UAS-FLP.D}JD1, P{Ubip63E(FRT.STOP)Stinger}9F6/CyO* to visualize endogenous *Drosophila Nlg3* expression [61]. Adult brains were dissected in 1x phosphate buffer saline (PBS) and fixed with 4% paraformaldehyde (PFA) for 20 minutes at room temperature and washed three times in 1% Triton/PBS (1% PBST). Samples were incubated overnight at 4°C with mouse anti-Repo [Developmental Studies Hybridoma Bank (DSHB, University of Iowa): 8D12, 1:200] and rat anti-Elav (DSHB: 7E8A10, 1:200) diluted in PBS/1% Triton/5% normal donkey serum (NDS). Following primary antibody staining, samples were washed three times with 1% PBST and incubated with anti-mouse-405 or anti-rat-405 (Jackson ImmunoResearch #715-475-151 and #712-475-153, 1:200) for two hours at room temperature, then mounted in Vectashield mounting media (Vector Laboratories #H-1000-10). Images were acquired using a Z-stack maximal projection on a LSM710 confocal microscope (Carl Zeiss) with a 10X objective lens. Images were processed using ImageJ software.

### *Drosophila* neuromuscular junction morphology

Muscle wall of third instar larvae (muscle 6/7 of segment A4) and abdominal body wall of 7-day-old adult flies were dissected on sylgard plates in 1X PBS and fixed with 4% PFA for 20 minutes. Samples were washed three times with 1% PBST and incubated overnight at 4°C with primary antibodies diluted in 1% PBST and 5% NDS. After three additional 1% PBST washes, samples were incubated with secondary antibodies for two hours at room temperature, mounted on Vectashield mounting media and imaged using a Z-stack maximal projection on a LSM710 confocal microscope with a 40X oil objective lens. Images were processed using ImageJ software. Primary antibodies: mouse anti-DLG (DSHB: 4F3, 1:50), mouse anti-BRP (DSHB: nc82, 1:25), and mouse anti-CSP (DSHB: 6D6, 1:25). Secondary antibodies: Alexa Fluor 488 AffiniPure Goat Anti-Horseradish Peroxidase (Jackson ImmunoResearch #123-545-021, 1:200), mouse anti-Cy3 (Jackson ImmunoResearch #715-165-151, 1:200), mouse anti-647 (Jackson ImmunoResearch #715-605-151).

### Electrophysiology recordings

Electrophysiological recordings were performed as described [91]. Wandering third instar larvae were dissected and recordings were obtained from muscle 6 of abdominal segments A3 and A4 with HL-3 solution containing 1mM Ca^2+^ (7.2 pH). HL-3 solution consisted of 70 mM NaCl, 5 mM KCl, 20 mM MgCl_2_, 10 mM NaHCO_3_, 5 mM trehalose, 115 mM sucralose, and 5 mM HEPES. Microelectrodes (10-20 MΩ) made with borosilicate glass were pulled using a micropipette puller (P-97; Sutter Instrument, California, USA) and filled with 3M KCl. Spontaneous miniature excitatory junction potentials (mEJPs) were recorded with an Axoclamp 900A amplifier (Molecular Devices, California, USA) in bridge mode, digitized with Digidata 1550 digitizer (Molecular Devices), collected with pClamp10.4 software (Molecular Devices), and analyzed using AxoGraph X software. mEJPs were recorded over a period of 120 seconds. Only cells with a resting potential between –50 to –80 mV and an input resistance > 4mΩ were used for analysis. All intracellular recordings were performed at room temperature.

### FM1-43 Uptake Assay

FM1-43 dye uptake assays were performed as previously described [92]. Wandering third instar larval body walls (muscle 6/7 of segment A4) were dissected on Sylgard plates in Ca^2+^ free HL-3 solution. To stimulate endocytosis, samples were incubated with 10 μm FM1-43 dye in HL-3 solution containing 90mM KCl and 1.5mM Ca^2+^ for 5 minutes then washed five times with Ca^2+^ free HL-3 solution. Imaging was performed on a LSM800 Airyscan confocal microscope (Carl Zeiss) with a 63x water objective lens and analyzed using ImageJ.

### Statistical Analysis

Statistical analysis was performed in Prism v10.6 (GraphPad). Data was tested for normality and variance. Normally distributed data where equal variance was violated were analyzed using Welches ANOVA followed by Dunnet’s T3 multiple comparison test, normally distributed data with equal variance was analyzed using one-way ANOVA followed by Sidak’s multiple comparison test, and non-normal data used Kruskal-Wallis with Dunn’s test. *P* < 0.05 was considered significant. All data is visualized as a bar chart (mean ± STD, including individual points). Sample sizes are described in the figure legends.

## Data Availability

All data generated or analyzed during this study are included in the manuscript and supporting files.

## Acknowledgements

We would like to thank the Bloomington *Drosophila* Stock Center, Vienna *Drosophila* research center, and Kyoto *Drosophila* stock center for providing *Drosophila* reagents, and the Developmental Studies Hybridoma Bank for providing antibodies. We would also like to thank Paul C. Marcogliese, Ph.D. for his help in making the *NLGN3^Ref^* transgene and Anne Simon, Ph.D. for generously providing the *Nlg3^Null^* fly lines.

## Contributions

RET performed most of the experiments with help from JCA on the behavioral experiments, SBH on the electrophysiology, SS on the FM1-43 dye loading, and SVJ on the immunohistochemistry. RET and OK generated the reagents. RET, SY, and MFW designed the project. RET and MFW wrote the paper with input from all other authors.

## Fig. legends

**Figure S1: *NLGN3* variants are well conserved across other *NLGN* family members. A)** Evolutionary conservation of amino acids across other human NLGN family members. **B, E)** *Nlg3^TG4^* driving a G-trace reporter line was used to identify the developmental expression of *Nlg3*. **C)** anti-Elav staining labels neurons. **D)** Merging the expression of *Nlg3* and Elav-positive cells identifies that *Nlg3* is expressed in neurons during development. **F)** anti-Repo staining labels glial cells. **G)** Merging *Nlg3* and Repo-positive cells shows some overlap between *Nlg3* and glia during development. Scale bar: 100 μm.

**Figure S2: Overexpression of the p.R175W variant alters sleep behaviors. A)** Quantification of the activity per waking minute (n≥19 flies; Welch’s ANOVA with Dunnett’s T3 multiple comparison test, ns *P* > 0.05, * *P* ≤ 0.05, ** *P* ≤ 0.01, **** *P* ≤ 0.0001).

**B)** Quantification of the total activity measured by video recording adult flies (n≥20 flies). **C)** Quantification of total activity in *NLGN3* overexpression models using a neuronal GAL4 driver (n≥30 flies). **D)** Quantification of activity per waking minute in *NLGN3* overexpression models (n≥30 flies). **E)** Quantification of sleep duration during lights on (n≥30 flies, Welch’s ANOVA with Dunnett’s T3 multiple comparison test, ** *P* ≤ 0.01, **** *P* ≤ 0.0001). **F)** Quantification of sleep duration during lights off (n≥30 flies). **G)** Quantification of sleep latency (n≥30 flies). **H)** Quantification of average sleep bout number during lights off (n≥30 flies, Welch’s ANOVA with Dunnett’s T3 multiple comparison test, ns *P* > 0.05, *** *P* ≤ 0.0001). **I)** Quantification of average sleep bout length during lights off (n≥30 flies). Kruskal Wallis with Dunn’s multiple comparison test, for B-D, F, G, and I; ns *P* > 0.05, * *P* ≤ 0.05, ** *P* ≤ 0.01, *** *P* ≤ 0.001, **** *P* ≤ 0.0001.

**Figure S3: *NLGN3* variants do not impair branching.** Quantification of the number of branches in larval neuromuscular junctions at muscle 6/7 of abdominal segment A4 from *y^1^ w*, Nlg3^TG4/Null^, Nlg3^Null/Null^, NLGN3^Ref^; Nlg3^TG4/Null^, NLGN3^R175W^; Nlg3^TG4/Null^, NLGN3^R451C^; Nlg3^TG4/Null^*, and *NLGN3^R597W^; Nlg3^TG4/Null^* larvae (n≥22 larvae, Kruskal-Wallis test; *P* = 0.0929).

**Figure S4: Overexpressing *NLGN3* variants with neuronal drivers alter synapse morphology. A)** Representative images of type 1B boutons at muscle 6/7 of abdominal segment A4 from overexpressing *LacZ*, *NLGN3^Ref^, NLGN3^R175W^, NLGN3^R451C^*, and *NLGN3^R597W^* using a *Nlg3^TG4^* driver. **B)** Quantification of bouton number (n≥16 larvae). **C)** Representative images of type 1B boutons at muscle 6/7 of abdominal segment A4 from overexpressing *LacZ*, *NLGN3^Ref^, NLGN3^R175W^, NLGN3^R451C^*, and *NLGN3^R597W^* using a neuronal driver (Elav-GAL4). **D)** Quantification of bouton number (n≥17 larvae). **E)** Representative images of type 1B boutons at muscle 6/7 of abdominal segment A4 from overexpressing *LacZ*, *NLGN3^Ref^, NLGN3^R175W^, NLGN3^R451C^*, and *NLGN3^R597W^* using a motor neuron (D42-GAL4) driver. **F)** Quantification of bouton number (n≥20 larvae). **G)** Representative images of type 1B boutons at muscle 6/7 of abdominal segment A4 from overexpressing *LacZ*, *NLGN3^Ref^, NLGN3^R175W^, NLGN3^R451C^*, and *NLGN3^R597W^* using a muscle driver (Mef2-GAL4). **H)** Quantification of bouton number (n≥19 larvae). One-way ANOVA with Sidak’s multiple comparison test, ns *P* > 0.05,* *P* ≤ 0.05, ** *P* ≤ 0.01, **** *P* ≤ 0.0001.

**Figure. S5: Synaptic architecture is dysregulated post development. A-G)** Representative images of 7-day-old adult abdominal neuromuscular junctions (NMJ) from *y*^1^ *w*\**, Nlg3^TG4/Null^, Nlg3^Null/Null^, NLGN3^Ref^; Nlg3^TG4/Null^, NLGN3^R175W^; Nlg3^TG4/Null^, NLGN3^R451C^; Nlg3^TG4/Null^*, and *NLGN3^R597W^; Nlg3^TG4/Null^* flies stained with anti-CSP, a presynaptic marker. Scale bar: 10 μm. **H)** Quantification of bouton number in an adult NMJ (n≥20 flies). One-way ANOVA with Sidak’s multiple comparison test; ns *P* > 0.05, *** *P* ≤ 0.001, **** *P* ≤ 0.0001.

**Figure S6: *NLGN3* variants do not alter mEJP kinetics. A)** quantification of the amplitude of miniature excitatory junction potentials (mEJP) in *y^1^ w*, Nlg3^TG4/Null^, Nlg3^Null/Null^, NLGN3^Ref^; Nlg3^TG4/Null^, NLGN3^R175W^; Nlg3^TG4/Null^, NLGN3^R451C^; Nlg3^TG4/Null^,* and *NLGN3^R597W^; Nlg3^TG4/Null^* larvae (n≥11 larvae; Kruskal-Wallis test, ns *P* = 0.3050). **B)** Quantification of mEJP rise time (n≥12 larvae, Kruskal-Wallis test, ns *P* = 0.1965). **C)** Quantification of mEJP decay time (n≥9 larvae; Kruskal-Wallis test, ns *P* = 0.0664). **D)** Representative mEJP traces recorded from muscle 6 at abdominal segments A3 and A4 from *y w, Nlg3^TG4/Null^, Nlg3^Null/Null^, NLGN3^Ref^; Nlg3^TG4/Null^, NLGN3^R175W^; Nlg3^TG4/Null^*, *NLGN3^R451C^; Nlg3^TG4/Null^*, and *NLGN3^R597W^; Nlg3^TG4/Null^* larvae.

**Figure S7: *NLGN3* rescues endocytic defects. A-F)** Representative images of boutons from *y*^1^ *w*\*, *Nlg3^TG4/Null^, Nlg3^Null/Null^, NLGN3^Ref^; Nlg3^TG4/Null^, NLGN3^R175W^; Nlg3^TG4/Null^, NLGN3^R451C^; Nlg3^TG4/Null^*, and *NLGN3^R597W^; Nlg3^TG4/Null^* larvae loaded with FM1-43 dye. **H)** Quantification of the FM1-43 dye uptake measured by the intensity of the dye normalized to the size of the boutons (N≥6 larvae, n=5 boutons per larva), One-way ANOVA with Sidak’s multiple comparison test; ns *P* > 0.05, * *P* ≤ 0.05, * *P* ≤ 0.01, *** *P* ≤ 0.001. Scale bar: 10 μm.

